# The colon is an extra-pancreatic source of glucagon: demonstration in a mouse model of pancreatectomy

**DOI:** 10.1101/2024.12.23.630024

**Authors:** Anne-Charlotte Jarry, Johanne Le Beyec, André Bado, Jean-François Gautier, Jean-Pierre Riveline, Bertrand Blondeau, Ghislaine Guillemain, Maude Le Gall

## Abstract

Glucagon is considered as a pancreas-specific hormone. However, an extra-pancreatic source of glucagon, which secretion is inappropriately induced by oral glucose, has been suspected for years in patients with diabetes. In a mouse model of subtotal pancreatectomy, we recapitulated the abnormal secretion of glucagon in response to oral glucose observed in pancreatectomised patients and excluded the remaining pancreas as the source of glucagon. We showed that the colon can produce and secrete glucagon. Subtotal pancreatectomy resulted in an increased expression of PCSK2 protease in the colon that favors the maturation of proglucagon into glucagon over other glucagon-related peptides normally produced by enteroendocrine cells. These results identify a mechanism that could contribute to abnormal glucagon production and poor glycemic control in pancreatectomized diabetic subjects. Moreover, our findings may pave the road to new therapeutic strategies to counteract poor glucose control for patients with diabetes.

## Introduction

Glucose storage and mobilization is tightly controlled by pancreatic hormones glucagon and insulin secreted by pancreatic α and β cells respectively. When insulin production is absent or insufficient, glucose accumulates in the blood leading to hyperglycemia, the prognostic criteria of diabetes. Since elevated blood glucose associates with cardiovascular, renal and neural diseases, therapies to lower blood glucose have been developed and used for decades, mostly by providing exogenous insulin or improving insulin action. Yet, observations have led to propose that uncontrolled glucagon secretion may also be a crucial actor of the disturbed glucose homeostasis in diabetes. In fact, patients with diabetes exhibit abnormal glucagon secretion, *i.e.* stimulated by oral glucose but impaired after insulin-induced hypoglycemia, that could contribute to their poor glycemic control (Campbell and Drucker, 2015). Moreover, glucagon signaling blockade either pharmacologically or genetically prevents the development of diabetes in streptozotocin-treated mice (Johnson et al., 1982; Lee et al., 2011) although this result has been challenged lately (Damond et al., 2016; Neumann et al., 2016). Because of these inversed glucagon secretions in response to glucose, one could propose that deregulated glucagon secretions may not originate from pancreatic α cells but from other cells. Recently, Lund and collaborators reported increased glucagon levels in the plasma of pancreatectomized patients following an oral charge of glucose, establishing the existence of an extra-pancreatic source of glucagon with no formal identification (Lund et al., 2016). We also reported postprandial glucagon secretion in four patients with pancreatectomy-associated diabetes (Riveline et al., 2016). A 3-days treatment with somatostatin of one of those patients efficiently suppressed glucagon secretion and was associated with glucose tolerance improvement (Riveline et al., 2016). In all those patients, secretion of glucagon is induced by oral glucose and resembles that of the gut hormone glucagon-like peptide 1 (GLP-1), which is matured from the same prohormone than glucagon: proglucagon (Rouillé et al., 1995; Sandoval and D’Alessio, 2015). Action of different pro-hormone convertases leads to differential proglucagon cleavage: in pancreatic α-cells prohormone convertase 2 (PCSK2) enzyme, together with its regulator SCG5 (secretogranin 5), allow the formation of glucagon, while in enteroendocrine cells PCSK1/3 enzyme processes proglucagon mainly into glicentin, GLP-1 and GLP-2 (Rouillé et al., 1995). It is now clearly recognized that some pancreatic α-cells co-produce glucagon and GLP-1 (Chambers et al., 2017; Whalley et al., 2011) and numerous studies on mouse models of insulin-resistance or diabetes describe an increase in the number of α-cells expressing PCSK1/3 and GLP-1 (Kilimnik et al., 2010; Whalley et al., 2011). We hypothesized that in specific conditions, *e.g.* diabetes subsequent to pancreatectomy, some enteroendocrine cells could mature proglucagon into glucagon, in addition or not to GLP-1, and secrete glucagon in response to oral glucose, providing therefore a pathologic extra-pancreatic source of glucagon.

## Results

### Glucagonemia in a mouse model of subtotal pancreatectomy

To better understand glucagon production after pancreatectomy, we set up a mouse model of subtotal pancreatectomy. Pancreatic lobes were resected, preserving some pancreatic parenchyma between the duodenum and the pancreatic duct but allowing an 84% pancreatic resection with only 7% of preserved islets (figure 1A-C). Pancreatectomized (Px) mice developed hyperglycaemia, ketonemia and glycosuria as soon as 2 days post-surgery (figure 1D-F). Maximal hyperglycaemia occurred after 7 days and was correlated with the amount of resected pancreas (figure 1G).

**Figure 1.**
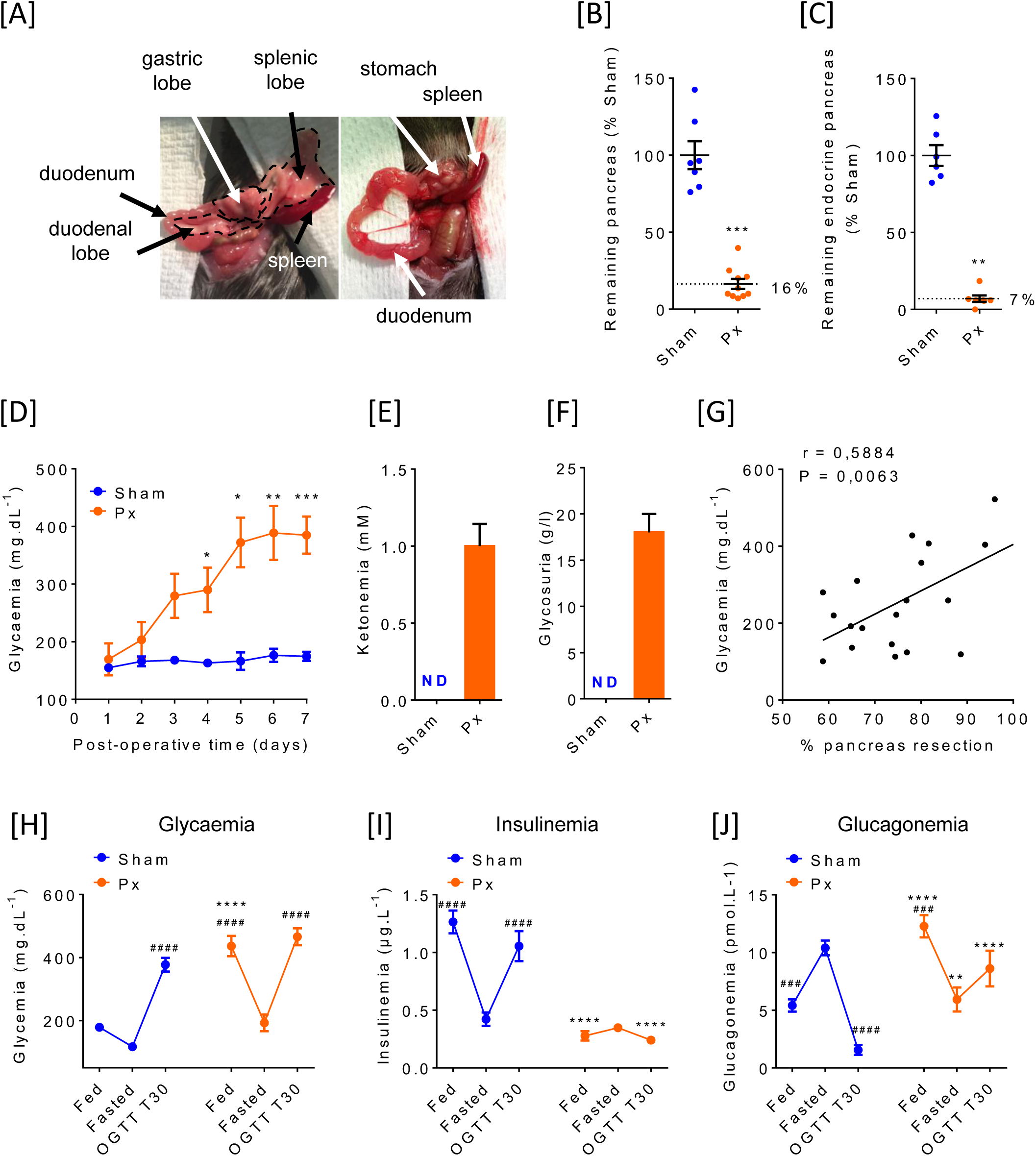
A murine model of subtotal pancreatectomy that recapitulates dysregulated glucagon plasma levels. (A) Mouse upper gut before (left panel) and after (right panel) pancreatectomy. (B) Weight of remaining pancreas 2 weeks post-pancreatectomy (Px) (n=10, orange circles) compared to sham mice (n=7, blue circles) represented as percentage of sham weight mean. (C) Number of pancreatic islets isolated from pancreas of sham (n=6, blue circles) *versus* Px (n=7, orange circles) represented as percentage of mean of sham islet number. (D) Blood glucose level in fed sham (n=7, blue circles) *versus* Px (n=9, orange circles) mice. Data are presented as mean ± SEM. (E) Ketonemia and (F) glycosuria in sham (n=6) *versus* Px (n=6) mice one week after surgery. ND not detectable. Data are presented as mean ± SEM. (G) Correlation between percentage of remaining pancreas and 7-day post-surgery fed glycaemia (n=19). (H) Glycaemia, (I) insulinemia and (J) glucagonemia of sham (n=5-7, blue circles) *versus* Px (n=5-11, orange circles) mice one week post-surgery in fed or fasted state and 30min following an oral load of glucose. Data are presented as mean ± SEM. *p<0.05, **p<0.01, ***p<0.001 ****p<0.0001 Px versus sham in similar condition. ^##^p<0.01, ^###^p<0.001, ^####^p<0.0001 Fed or OGTT T30 versus Fasted in sham or in Px animals.

Px mice displayed hyperglycaemia at fed states compared to sham animals (figure 1H). Fed sham animals displayed high insulin levels that decreased with fast and increased when a glucose load was given (figure 1I). In contrast, Px mice presented very low levels of insulin in all states (figure 1I), validating that subtotal pancreatectomy led to undetectable blood insulin. Plasma glucagon levels increased in fasted sham animals and decreased in response to an oral glucose load. In opposition, glucagon levels in Px mice were high in the fed state, decreased with fasting and increased in response to an oral glucose load compared to sham animals (figure 1J). Thus, this mouse model of subtotal pancreatectomy recapitulates the hormonal profile of pancreatectomized patients (Lund et al., 2016; Riveline et al., 2016).

### Remaining pancreas is not the source of glucagon in Px mice

Since murine subtotal pancreatectomy was previously studied as a model of endocrine cell regeneration (Bonner-Weir et al., 1993), we examined the remnant pancreas histology and endocrine cell composition. After subtotal pancreatectomy, the remaining pancreas underwent morphological modifications with apparition of fibrosis (figure 2A-B), characteristic of pancreatitis. We evaluated endocrine cell morphometry and measured insulin and glucagon production in the remaining pancreas immediately after pancreatectomy (Px0) and up to 2 weeks later (Px15). Two weeks after pancreatectomy, β-cell mass increased in the remaining pancreas, as previously reported (Bonner-Weir et al., 1993; Dor et al., 2004; Xu et al., 2008), while α-cell mass remained unchanged compared to the remaining pancreas at day 0, *i.e*. just after pancreatectomy (Figure 2C and 2D). Pancreas has been shown to contain 0.86 pmol.mg^−1^ of glucagon in control mice (Thorel et al., 2011). Pancreatic remnants contained 0.085 pmol.mg^−1^ of glucagon just after pancreatectomy (Px0) and 0.011 pmol.mg^−1^ 7 days later (Px 7) (Figure 2E). In pancreatic islets isolated from sham animals, an increase in insulin secretion and a decrease in glucagon secretion were detected in response to low concentration compare to high glucose concentration (figure 2F-G). However, no glucagon secretion by isolated islets from Px mice could be detected in response to either high or low glucose concentration (figure 2G). Finally, insulinemia (figure 2H) but not glucagonemia (figure 2I) was correlated with the percentage of remaining pancreas. All these arguments led us to propose that the remaining pancreas was not the source of glucagon secreted in the fed state or in response to an oral glucose load observed 7 days after pancreatectomy.

**Figure 2.**
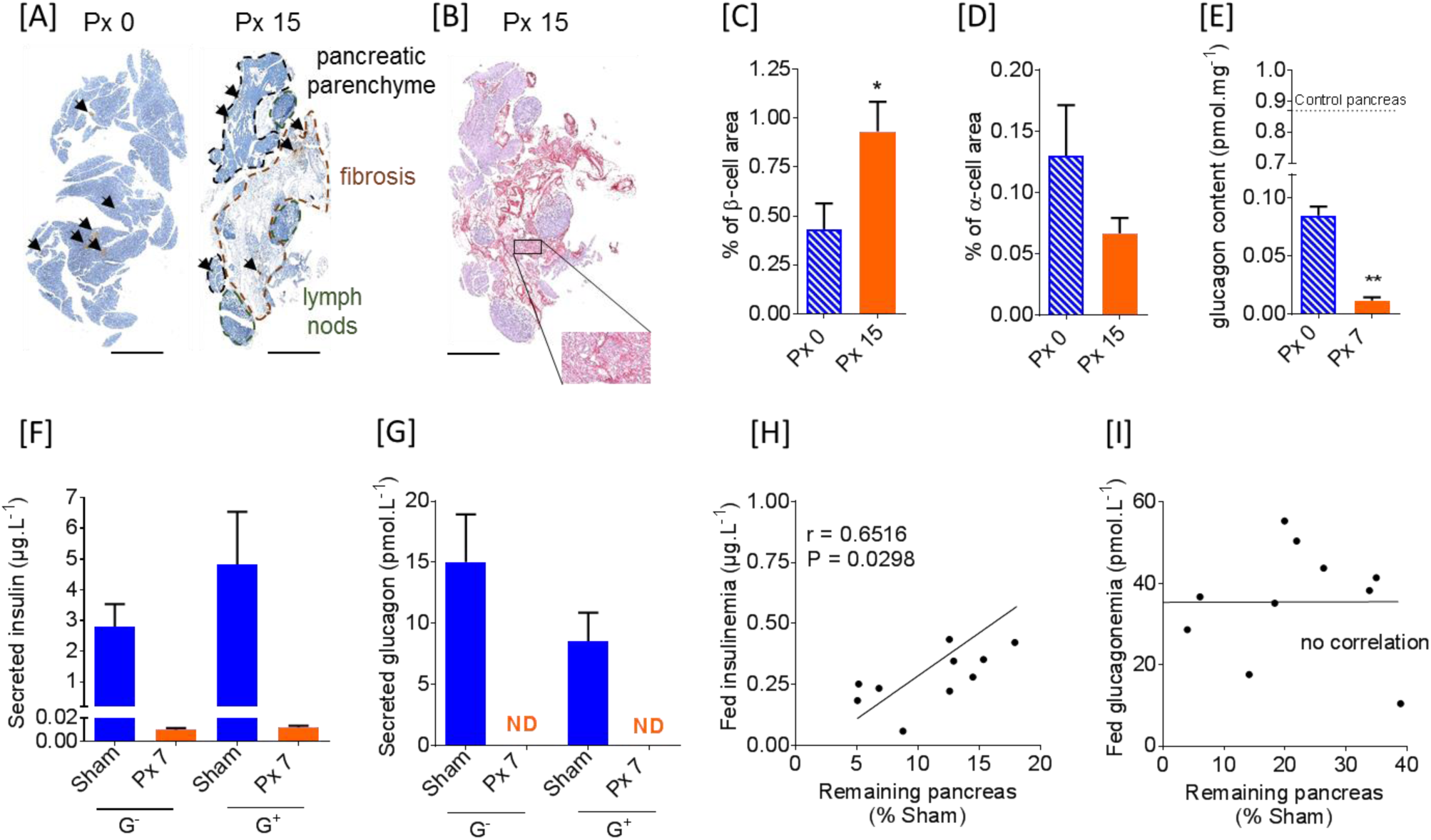
The remaining pancreas is not the source of glucagon secretion in Px mice. (A) and (B) are pictures of representative sections of the remaining part of the pancreas dissected immediately (Px0) or 2 weeks (Px15) after pancreatectomy. Scale bar 150 µm. (A) Immunohistochemistry for insulin (brown) followed by hematoxylin staining. Arrows point to cluster of β cells. (B) Picrosirius red staining on a section of the remaining pancreas 2 weeks (Px 15) after pancreatectomy. Note the important fibrosis observed 15 days post-surgery. (C) β- and (D) α-cell area expressed as % of total tissue area in the remaining pancreas of Px mice sampled immediately (Px0, n=4-5, hatched blue bars) or 2 weeks (Px15, n=6, orange bars) post-surgery. (E) Glucagon content in the remaining pancreas per mg of tissue in Px mice sampled immediately (Px0, n=5, hatched blue bars) or 7 days (Px7, n=6, orange bars) post-surgery. (C-E) Data are presented as mean ± SEM *p<0.05, **p<0.01 Px7 versus Px0 (F-G) Insulin (F) and Glucagon (G) secreted by 15 pancreatic islets isolated from sham (n=6, blue bars) and Px (n=12, orange bars) mice 7 days post-surgery in response to 1mM (G-) or 11mM (G+) glucose. Data are presented as mean ± SEM. ND not detectable. (H-I) Correlations the between percentage of remaining pancreas and fed insulinemia (H) or fed glucagonemia (I) 7 days post-surgery (n=10).

### Colon produces and secretes glucagon in Px mice

As *proglucagon* gene is expressed in pancreatic α-cells but also in enteroendocrine cells scattered along the gastrointestinal tract, we investigated the possibility of glucagon production by enteroendocrine cells. First, we measured proglucagon expression in the different parts of the gut. As already reported in the literature (Sandoval and D’Alessio, 2015), we found that proglucagon expression was higher in the colon than in other parts of the gut of sham animals (figure 3A). mRNA coding PCSK1/3 – responsible for the maturation of proglucagon into GLP-1 in enteroendocrine cells – is expressed throughout the gut (figure 3B). However, mRNA coding PCSK2 – responsible for the maturation into glucagon – was only detected in the colon in sham and Px mice and colonic PCSK2 mRNA expression was dramatically increased in Px compared to sham mice (figure 3C). In parallel, we observed that mRNA coding SCG5 (a regulator of PCSK2) was also expressed in the colon of mice without any difference between sham and Px mice (data not shown). Since *proglucagon*, *Pcsk1/3* and *Pcsk2* are all expressed in the colon, we focused our interest on this distal part of the gut. We observed GLP-1 as well as glucagon-producing cells by immunofluorescence on colonic sections of sham and Px mice (figure 3D). Glucagon-expressing cells had a pear-shape similar to enteroendocrine cells and in some cells, glucagon co-localized with GLP-1 (figure 3E). After pancreatectomy, the ratio of glucagon-over GLP-1-positive cells doubled (figure 3F) in Px compared to sham mice, together with an increased ratio of *Pcks2* over *Pcsk1/3* mRNA levels (figure 3G). Accordingly, glucagon protein content presented a 4.1-fold increase in the colon of Px mice versus sham animals, whereas it was not different in the other gastrointestinal segments (figure 3H). The increased colonic content of glucagon in Px mice was associated with an increased capacity of the colon to secrete glucagon measured *ex vivo* (figure 3I). Finally, colonic secretion of glucagon *ex vivo* was correlated with fed glucagonemia *in vivo* (figure 3J) strongly suggesting that the colon could be a source of deregulated secretion of glucagon after pancreatectomy.

**Figure 3:**
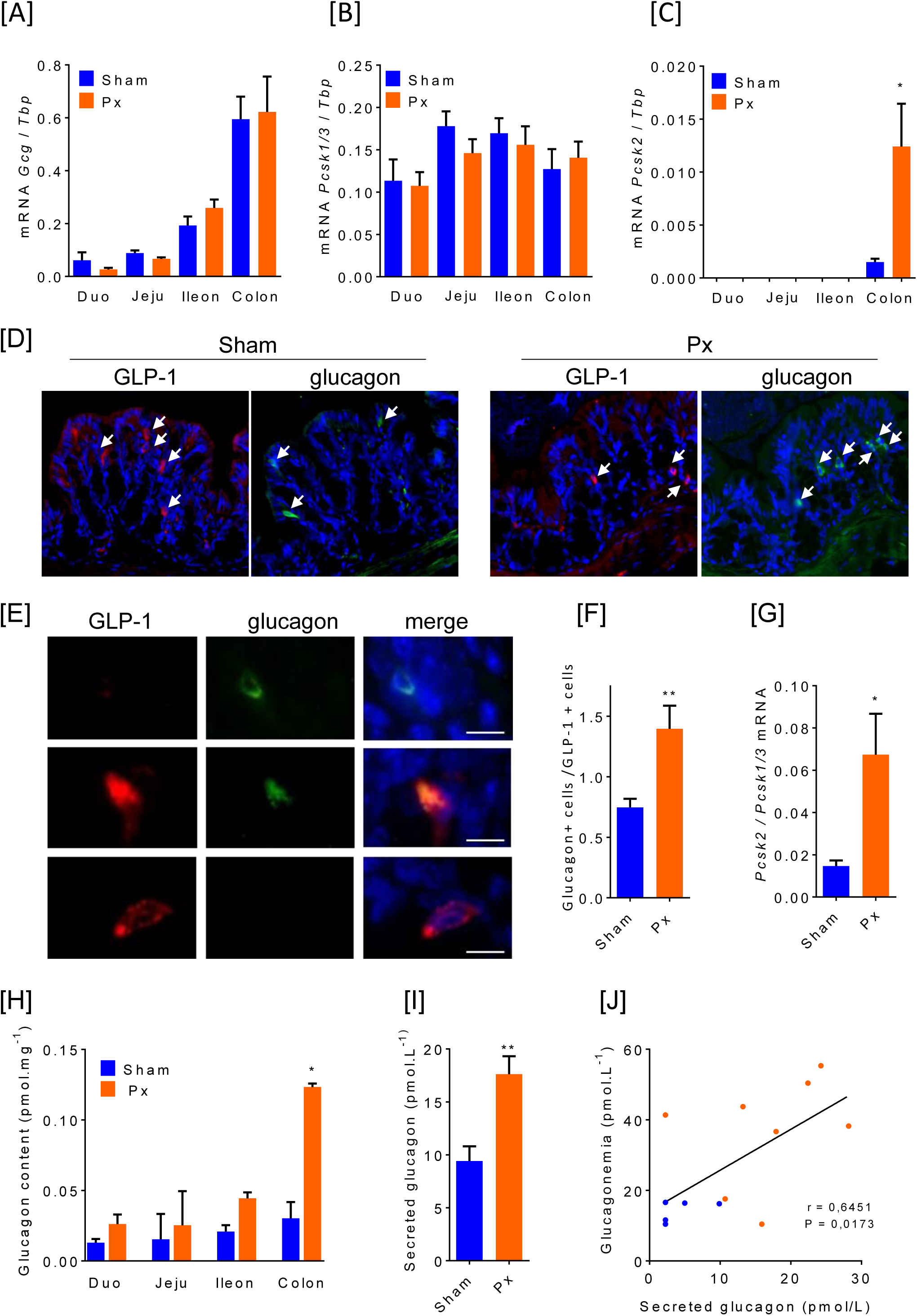
Adaptation of the gut after subtotal pancreatectomy results in an increased glucagon production and secretion by the colon. (A-C) mRNA levels of *Gcg* (A), *Pcsk1/3* (B), *Pcsk2* (C) normalized to the reference gene *Tbp* along the gastro-intestinal tract of sham (n=6, blue bars) and Px mice (n=8, orange bars). (D) Immunofluorescence of GLP-1 (red), glucagon (green) and nuclei (blue) on sequential sections of colon from sham or Px mice one week post-surgery. Scale bar = 50µm. (E) Co-immunofluorescence of GLP-1 (red), glucagon (green) and nuclei (blue) on colonic sections of Px mice one week post-surgery. Scale bar = 10µm. (F) Ratio of GLP-1-over glucagon-immunoreactive cells in the colon of sham (n=9, blue bars) and Px (n=8, orange bars) mice. (G) Ratio of colonic *Pcsk2* over *Pcsk1*/3 mRNA levels in sham (n=5, blue bars) and Px (n=8, orange bars) mice. (H) Glucagon content along the gastrointestinal tract of sham (n=5-7, blue bars) and Px (n=6-8, orange bars) mice. (I) *Ex vivo* secretion of glucagon by a colonic portion (1 cm of length) of sham (n=6, blue bars) and Px (n=12, orange bars) mice. (K) Correlation between *in vivo* fed glucagonemia and *ex vivo* colonic glucagon secretion of sham (n=5, blue circles) and Px mice (n=8, orange circles). (A, B, C, F, G, H I) Data are presented as mean ± SEM. *p<0.05, **p<0.01 Px *versus* sham.

## Discussion

An extra-pancreatic source of glucagon responsible of brittle fluctuations of glycaemia reported in diabetic patients has been alleged and debated for more than 50 years (Unger et al., 1966), but the recent demonstration that glucagon is present in patients with total pancreatectomy reopened the debate (Lund et al., 2016; Riveline et al., 2016). As proglucagon is produced not only in the pancreas but also in the gut, we hypothesized that in specific metabolic conditions like pancreatectomy, proglucagon may be matured into glucagon in enteroendocrine cells and secreted in response to an oral load of glucose.

Subtotal pancreatectomy in mice is usually performed to assess the ability of β cells to regenerate (Bonner-Weir et al., 1983). Following pancreatic injury, new β cells appear either from replication of pre-existing β cells (Xu et al., 2008) or from differentiation of ductal precursors (Dor et al., 2004). These pancreatic adaptations take place no longer than 1-day post-surgery (Bonner-Weir et al., 1993). However, to our knowledge, no study reported α-cell adaptation to subtotal pancreatectomy in mice. Two weeks post-surgery, we observed a slight tendency for a decreased α-cell area suggesting that, contrary to β cells, α cells do not regenerate following subtotal pancreatectomy. Ability of α cells to convert into β cells has been documented in a diabetic state resulting from a nearly absolute β cell ablation (Thorel et al., 2011), and could occur after subtotal pancreatectomy. As we assay α-cell area by glucagon staining, one can also hypothesis that maturation of pancreatic proglucagon into glucagon is altered after pancreatectomy as some studies report a switch toward production of GLP-1 in α cells in other pathological conditions (Whalley et al., 2011). Pancreatectomy is associated with the settlement of fibrosis characteristic of pancreatitis which is associated with IL6 production (Pooran et al., 2003). IL6 is a pro-inflammatory cytokine that has been shown to stimulate GLP-1 production by α cells *ex vivo* (Ellingsgaard et al., 2011; Timper et al., 2016). The inflammatory state in the remaining pancreas after pancreatectomy may favour GLP-1 production rather that glucagon production by α cells.

The absence of α-cell regeneration, associated with the decrease in pancreatic glucagon production and secretion following pancreatectomy, led us to propose that the pancreas was not the source of fed hyperglucagonemia and glucagon secretion reported in response to an oral load of glucose. Conversely, we demonstrated that glucagon production and secretion by the distal gut increased following pancreatectomy. Colonic glucagon secretion *ex vivo* was correlated to fed glucagonemia *in vivo* supporting the gut as the source of hyperglucagonemia in pancreatectomised mice. Conflicting results concerning glucagon production by the gastro-intestinal tract probably reflected the lack of specificity of many glucagon assays (Tanjoh et al., 2003; Wewer Albrechtsen et al., 2014). Proglucagon is processed into a wide variety of peptides, some of which share common sequences with glucagon _(33-61)_. Glucagon assay using antibodies directed against the N-terminal fraction of glucagon also detect oxyntomodulin _(33-69)_ and those using antibodies directed against the C-terminal fraction of glucagon detect miniglucagon _(51-61)_ but also high levels of oxyntomodulin _(33-69)_, and glicentin _(1-69)_. A recent study report 18 new peptides derived from proglucagon that share common sequences with glucagon (Wewer Albrechtsen et al., 2017). In the present study, we used a highly specific glucagon assay that uses two antibodies targeting both N- and C-terminal ends of glucagon reducing cross-contaminations with all those proglucagon-derived products (Roberts et al., 2018; Wewer Albrechtsen et al., 2014). Additionally, our immunostaining experiments with 2 different anti-glucagon antibodies confirm its presence in colonic cells that co-express or not GLP-1. Still, we cannot formally exclude that we did not detect proglucagon variant PG 1-61, a peptide sharing common sequences with glucagon that is highly expressed in diabetic condition (Wewer Albrechtsen et al., 2017). However, PG 1-61 possesses gluco-regulatory effects through activation of the glucagon receptor, thus the hyperglucagonemia may therefore, not only reflect hypersecretion of fully processed glucagon, but also of PG 1-61.

Demonstrating the overexpression of PCK2 over PCSK1/3 in the colon, we were able to dissect some of the molecular mechanisms that allow the production of glucagon by the colon. Colocalisation of GLP1 and glucagon suggests that the production of glucagon by the gut may originate from enteroendocrine L-cells that adapted their phenotype and express PCK2 over PCSK1/3 and produce glucagon in addition to GLP-1. Similar changes have been recently reported for other enterohormones in short bowel enteroendocrine cells along the crypto-villus axis (Beumer et al., 2018). This plasticity reflected by the relative expression of PCK2 over PCSK1/3 and activated upon metabolic alterations has already been reported in α cells (Kilimnik et al., 2010; O’Malley et al., 2014; Whalley et al., 2011) and could be a common feature shared with L cells. However, it cannot be excluded that in the distal gut, glucagon-producing cells arise from the differentiation of new precursors located in colonic crypts.

In accordance with our results, Jorsal *et al*. reported in human gut that *PCSK1/3* and *PCSK2* expression are increased patients with type 2 diabetes compared with healthy individuals (Jorsal et al., 2018)). They proposed that this increased of *PCSK2* expression in the gut could be responsible for the hyperglucagonemia observed in patients with type 2 diabetes.

Our demonstration of the existence of colonic glucagon in pancreatectomized mice validates observations already made in pancreatectomized patients (Lund et al., 2016; Riveline et al., 2016) and is the first step toward the understanding of the brittle fluctuations of glycaemia reported in these patients. Whether the colon also overproduces glucagon in type 1 and type 2 diabetes is of particular interest since it may contribute to impaired glycaemic control. Our murine model will help decipher the signal(s) responsible for this overproduction and identify pharmacological targets to prevent colonic glucagon production that will be of interest in the treatment of patients with diabetes.

## Acknowledgment

We are grateful to Prof S. Bonner-Weir for providing help in setting up subtotal pancreatectomy in mice, O. Thibaudeau (Anapathology Platform Bichat Hospital) and Prof A. Couvelard (Anapathology Service Bichat Hospital) for help with IHC acquisition and interpretation, C. Klein (Cordelier Research Center imagery Platform) for help with image acquisition. We also thank A. Gélineau, C. Moulineuf, A. Goncalves, A. Mevel and S. Bouanane for their technical help and Prof. S Ledoux and Prof E. Larger for discussions and reviewing of the manuscript.

## Methods

All animal experiments described in this study were conducted in compliance with the European Community guidelines and were approved by the Institutional Animal Care and Use Committee (*N° 11256* Comité d’Ethique Paris-Nord N°121). Animals were acclimated to the facility for at least one week before studies were initiated. They were maintained on a 12h light-dark cycle with free access to water and chow diet (Altromin1324, Genestil, France). All efforts were made to minimize the number of animals used and their suffering.

### Animal surgery

C57BL/6J male mice (2 to 3-month-old, Janvier, France) were either pancreatectomized (Px) or sham-operated. After anesthesia with isoflurane (Vetflurane, Virbac), a midline laparotomy was performed. The spleen and the stomach were isolated allowing resection of the pancreatic tail. The duodenal loop was isolated to resect pancreatic head. Pancreatic head and tail were weighted to assess the percentage of pancreatic resection as follow: (resected pancreas weight during surgery/body weight) / (sham pancreas weight at sacrifice/body weight) x100. Little pancreatic parenchyma between the pancreatic duct and the duodenum was conserved. In sham-operated mice, stomach and spleen were isolated and the pancreas was tweaked before being replaced in the abdominal cavity. In both cases, laparotomy was closed with 7.0 polypropylen suture in 2 layers. Fed weight and glycaemia were evaluated daily with AccuChek System (Roche Diagnostics) and expressed in mg.dL^−1^. Ketonemia was assessed with FreeStyle Optimum Neo (Siemens), glycosuria was measured with Multistix 8 SG (Siemens) one week post-surgery.

### Plasma insulin and glucagon measurements

5 days post-surgery, after a 4 hour-fast period and 30 minutes following a glucose oral load (2g/kg), blood (200 µL per mice) was collected by a retro-orbital puncture. Blood samples were centrifuged at 3000 g for 10 minutes and plasma were immediately stored at −20°C until subsequent analyses. Insulinemia and glucagonemia were measured using a Mouse Ultrasensitive Insulin ELISA kit (Alpco) and a Glucagon 10μL ELISA kit (Mercodia) respectively, according to manufacturers’ instructions.

### Measurement of glucagon tissue contents

Remaining pancreas were collected immediately (Px 0) and 1 week (Px) after pancreatectomy. In sham and pancreatectomized (Px) mice, duodenal, jejunal, ileal, and colonic mucosa were scrapped and isolated 1 week after surgery. Samples were homogenized in ethanol/acid (ethanol: sterile water: 12N HCl 75:24:1 v/v) solution (5 mL/g tissue). After an overnight incubation at 4°C, samples were centrifuged at 3000g for 20min at 4°C, supernatants were stored at −80°C for subsequent analyses. Tissue glucagon content was measured using Glucagon 10μL ELISA kit (Mercodia) according to manufacturers’ instructions. (Wewer Albrechtsen et al., 2014)

### Histology and immunochemistry

Remaining pancreas after pancreatectomy was collected immediately (Px 0), 1 week (Px 7) or 2 weeks (Px 15) post-surgery. To assess α- and β-cell areas, 3 µm sections of the all remnant pancreas were performed using a microtome as previously described. (Valtat et al., 2013) Sections were picked every 10 sections to perform glucagon or insulin immunostaining. Sections were incubated with anti-insulin (guinea-pig polyclonal Dako) or anti-glucagon (rabbit polyclonal, Diasorin) antibodies and then with secondary antibodies coupled to HRP. The HRP substrate DAB (Dako) was then used to stain α- or β-cells. The α- and β-cell areas were calculated as the ratio of pancreatic glucagon- or insulin-positive cell area to total tissue area of the entire section, as determined by computer-assisted measurements, using a Leica DMRB microscope equipped with a color video camera coupled to a Leica Q500IW PC computer, as previously described. (Valtat et al., 2013) For morphometric analyses, pancreatic sections were used to perform picrosirius red staining. Slides were scanned with an Aperio ScanScope CS System (Leica Microsystemes SAS).

Proximal colonic segments were sampled from sham and Px mice one week after surgery. Samples were fixed overnight in 4% paraformaldehyde in 10 mM PBS, pH 7.4, dehydrated and embedded in paraffin. Serial sections of 3µm from mouse colonic segments were used for immunostaining using GLP-1 (mouse monoclonal ab26278 Abcam) and glucagon (mouse monoclonal ab10988 Abcam) antibodies. In some experiments, GLP-1 (mouse monoclonal ab26278 Abcam) and glucagon (rabbit polyclonal, Diasorin) co-immunostaining were performed. After dewaxing and rehydrating, antigen retrieval was performed by pre-treatment with high temperature at pH 9. Colonic sections were then incubated with primary antibodies overnight at 4°C followed by secondary antibodies (mouse IgG ab150113 Abcam, rabbit IgG ab96891 Abcam) for 1 hour at room temperature. Slides were scanned with a Zeiss Axio Scan.Z1. Numbers of GLP-1 and glucagon positive cells were evaluated using Zeiss Zen 2 (Blue edition) software and normalized to the colonic mucosa surface to obtain GLP-1 and glucagon cells densities. Density ratio was calculated as follow: glucagon-positive cell density / GLP-1 positive cell density.

### Ex vivo glucagon secretion

Sham and pancreatectomized (Px) mice were anesthetized one week after surgery and pancreatic islets were isolated after injection of a 1 mg/mL collagenase (C7657, Sigma) solution in the bile duct. Injected pancreata were then incubated 20 min at 37°C and washed with 10% FBS HBSS. 15 islets per mouse were handpicked under a binocular microscope (Leica) and incubated in RPMI1640 (Invitrogen) + 10% fetal bovine serum (PAA). After 24 hours, islets were transferred in Krebs buffer containing 11 mM glucose (Sigma) for 1 hour. Then they were transferred in Krebs buffer containing 1 mM glucose for 1 hour. Supernatants were stored at −80°C until subsequent analyses.

1-cm long proximal colonic segments from anesthetized mice were sampled and incubated in Krebs buffer for 1 hour. Supernatant were stored at −80°C until subsequent analyses.

Supernatant glucagon concentration was measured using Glucagon 10 μL ELISA kit (Mercodia) according to manufacturers’ instructions.

### Reverse Transcription and Quantitative Real-Time Polymerase Chain Reaction

Total RNAs were extracted from pancreas or gastro-intestinal segments with TRIzol reagent (Invitrogen, Saint Aubin, France) according to manufacturer’s instructions. Complementary DNAs were obtained from one microgram of total RNA using the Verso cDNA Synthesis Kit (Thermo Scientific, Courtaboeuf, France). Real-time polymerase chain reaction was performed using the LightCycler480 system (Roche Diagnostics, Indianapolis) according to manufacturer’s instructions under the following conditions: 15-minute denaturation at 95°C, followed by 50 cycles of 10-second at 95°C, 45 seconds at 60°C, and 10 seconds at 72°C. Melting curves were generated for each reaction, from 55°C to 95°C at 0.11°C/s. Ct values of the target gene were normalized with 2 different reference genes (TBP and HPRT), which were chosen after comparison with numerous reference genes. The primers used in this study were designed using Roche assay design centre and synthetized by Eurogentec.

*Gcg Fwd: 5’-CAATGGCGACTTCTTCTGG-3’*

*Gcg Rev: 5’-CCAGTGATGTGAGTTCTTACTTGG-3’*

*Pcsk2 Fwd: 5’-GGCGTGTTTGCATTAGCTTT-3’*

*Pcsk2 Rev: 5’-GCACAGTCAGATGTTGCATGT-3’*

*Pcsk3 Fwd: 5’-TGGAGTTGCATATAATTCCAAAGTT-3’*

*Pcsk3 Rev: 5’-AGCCTCAATGGCATCAGTTAC-3’*

*Scg5 Fwd: 5’-ACCAGGCCATGAATCTTGTT-3’*

*Scg5 Rev: 5’-TCCTTAGGAATGTTGTCACCAG-3’*

### Statistical analyses

Results are expressed as mean ± SEM. Mann–Whitney U tests and Kruskal–Wallis tests followed by Dunn’s post-test were used to compare 2 groups and more than 2 groups respectively. 2-way ANOVA followed by Bonferroni post-test for multiple comparisons were used for time course analyses. P < 0.05 was considered significant.

## Abbreviations

GLP-1: Glucagon-like peptide 1
Px: pancreatectomized
Pcsk: prohormone convertase

## Author contributions

JFG and JPR initiated and discussed the project. ACJ, GG, JLB, BB, JPR and MLG designed the experiments. ACJ, GG, BB and MLG performed experiments and acquired data. ACJ, GG, JLB, BB, AB, JPR and MLG analyzed and interpreted data. ACJ and MLG wrote the manuscript with inputs from JLB, BB and GG.

